# Balancing energy supply during photosynthesis - a theoretical perspective

**DOI:** 10.1101/476846

**Authors:** Anna Matuszyńska, Nima P. Saadat, Oliver Ebenhöh

## Abstract

The photosynthetic electron transport chain (PETC) provides energy and redox equivalents for carbon fixation by the Calvin-Benson-Bassham (CBB) cycle. Both of these processes have been thoroughly investigated and the underlying molecular mechanisms are well known. However, it is far from understood by which mechanisms it is ensured that energy and redox supply by photosynthesis matches the demand of the downstream processes. Here, we deliver a theoretical analysis to quantitatively study the supply-demand regulation in photosynthesis. For this, we connect two previously developed models, one describing the PETC, originally developed to study non-photochemical quenching, and one providing a dynamic description of the photosynthetic carbon fixation in C3 plants, the CBB Cycle. The merged model explains how a tight regulation of supply and demand reactions leads to efficient carbon fixation. The model further illustrates that a *stand-by* mode is necessary in the dark to ensure that the carbon fixation cycle can be restarted after dark-light transitions, and it supports hypotheses, which reactions are responsible to generate such mode *in vivo*.

## 1 INTRODUCTION

Decades of multidisciplinary research of photosynthesis resulted in our today’s detailed understanding of the molecular, regulatory and functional mechanisms of light driven carbon fixation. Yet, still much is to uncover, especially in terms of identifying processes limiting photosynthetic productivity, calling for further basic research that may help redesigning photosynthesis [1, 2]. Historically, the process of photosynthesis has been divided into two parts. The so-called light reactions carried by the photosynthetic electron transport chain (PETC) convert light into chemical energy, supplying ATP and NADPH. This energy is next used to drive the carbon dioxide reduction and fixation in the processes known as the dark reactions. Thus, the metabolic light and dark reactions can be viewed as a molecular economy **supply-demand system** [3, 4, 5].

Despite this clear interdependence, these processes are often studied in isolation, permitting detailed and in-depth analysis of particular components at the cost of simplification of the preceding / following processes. Such separation is also reflected in theoretical research. Numerous approaches in the past decades aimed at translating the complexity of photosynthesis into a mathematical language, resulting in an impressive portfolio of kinetic models. The majority of these models focus either on the supply or on the demand side. Many classical models of the Calvin-Benson-Bassham (CBB) cycle, such as the biochemical models for C3 photosynthetic CO_2_ assimilation [6, 7, 8, 9, 10, 11, 12], made no attempt to model the processes of the PETC in any detail. Instead, they simplify the rate of electron transport supplying ATP and NADPH in often just one lumped reaction (e.g., non-rectangular hyperbola as a function of absorbed irradiance in [13]), or even kept them as a constant (NADPH in [9]). Likewise, many models of the PETC made no attempt to include details of the energy consuming reactions and describe ATP and NADPH demand by simple lumped reactions. Such an understandable simplification resulted from the fact that these models were created to study a specific light harvesting mechanisms, such as state transitions [14], non-photochemical quenching (NPQ) [15, 16, 17] or the role of antenna complexes in photosynthetic productivity [18].

The purpose of this study is to understand the interactions and interdependencies of the PETC and the carbon fixation cycle, with a focus on investigating the supply-demand control of photosynthesis. For this, we require a model that contains both processes but is simplified enough to perform a rigorous and systematic Metabolic Control Analysis (MCA) to derive general conclusions. Noteworthy, there exist a few successful attempts to include both electron transport and carbon assimilation processes into a unified mathematical framework. The model proposed by Laisk *et al.* [19] provides a solid summary of our knowledge on photosynthesis. The emphasis of the model structure was on including the electron transport through photosystems (PSII and PSI), while providing a detailed description of the down-stream metabolism. As a result, the model can represent steady state photosynthesis and chlorophyll fluorescence, but was insufficient to reproduce the dark-light induction of photosynthesis, a property that is critical in the context of our supply-demand research. A subsequent model of e-photosynthesis by Zhu *et al.* [20] is a comprehensive description of the process, that includes “as many photosynthesis-related reactions as possible”. Due to its complexity, employing the e-photosynthesis model [20] for a systematic supply-demand analysis is challenging. Moreover the highly detailed description of the molecular processes included in the model makes it hard to draw conclusions of general validity. Finally, Morales *et al.* [21] developed recently a thorough model of the PETC, including all relevant processes at the chloroplast and leaf level. Nevertheless, since the emphasis of this model was on the PETC regulation, the CBB cycle has been simplified into two steps. This imbalance in the levels of detail describing the two sub-processes is the main reason why we decided against using it.

Therefore, we have developed a new model that contains the key components of both subsystems, yet is simple enough to allow for systematic investigations. Using the quantitative theory of MCA, that investigates the effect of the perturbation of single processes on the overall stationary behaviour of the complete system [22, 23, 24], and metabolic supply-demand analysis [3] we aim at providing quantitative insight into the role of various photosynthetic processes in the regulation of their tight interdependence. The model has been constructed by merging a model of the PETC, originally designed to study photoprotective mechanisms [14, 17], with a kinetic model of C3 carbon fixation [9, 10]. We demonstrate that coupling these two models into a connected supply-demand system is possible, but far from trivial, and results in new emergent properties. We show that different light conditions and light protocols have effects on the stability of the carbon fixation cycle. We illustrate the need for a *stand-by* mode in the dark to ensure the restart of the carbon fixation cycle after dark-light transitions. Finally, we hint at new roles for long known reactions that keep carbon fixation intact and regulate its efficiency.

## 2 THE MODEL

We are presenting here the result of our exercise of merging together two independent, previously developed kinetic models of photosynthesis, both based on ordinary differential equations (ODEs). The first model describes the primary photosynthetic reactions through the PETC, leading to the production of ATP and NADPH. The CBB cycle is considered as the main consumer of the produced energy and reducing equivalents. Therefore in this model, the downstream metabolism has been simplified to two consuming reactions governed by mass action kinetics. It has been developed based on our previous work: the core model of the PETC by Ebenhöh *et al.* [14] and the model of high-energy dependent quenching in higher plants developed by Matuszyńska *et al.* [17]. Using this model, we are able to compute the fluorescence emission under various light protocols, monitor the redox state of the thylakoid and the rate of ATP and NADPH synthesis.

The second model is the Poolman [10] implementation of the carbon fixation model by Pettersson and Ryde-Pettersson [9], reproduced in our Institute using the modelbase software [25]. In contrast to the original model [9], in the Poolman representation the rapid equilibrium assumptions were not solved explicitly, but instead approximated by mass-action kinetics with very large rate constants. Solving the system of ODEs allows computation of different carbon fixation rates and reaction activities at varying concentrations of external orthophosphate. In the original model, the input of the ETC has been simplified by a single lumped reaction of ATP synthesis (*v*_16_ in [9]), whilst NADPH has been kept as a constant parameter.

### 2.1 Included processes and the stoichiometry

The model, schematically captured in Figure 1, comprises of 35 reaction rates and follows the dynamics of 24 independent state variables (see Supplement for a full list of reaction rates and ODEs). In addition, we compute a number of values such as emitted fluorescence or variables derived from conserved quantities. Light is considered as an external, time-dependent variable. Since the focus of this model is to study basic system properties, such as the response to relative changes in the light intensity, we did not calibrate our simulations to experimentally measured light intensities. Therefore, in this work light is expressed in µ mol photons per m^2^ and second and reflects the quantity of light efficiently used, but the utilization conversion factor to the Photon Flux Density (PFD) of the incident light is unknown.

**FIGURE 1.**
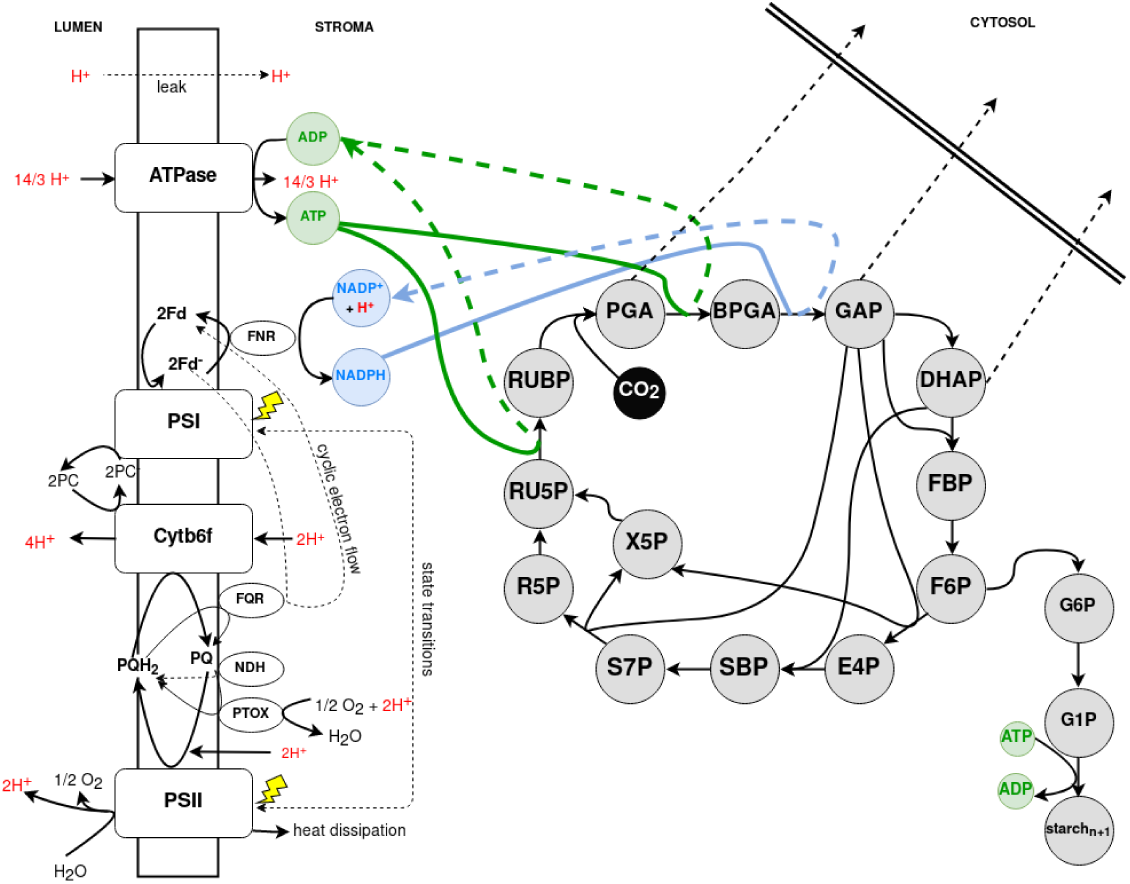
Schematic representation of the photosynthetic processes described by our *merged* mathematical model. The reactions take place in two compartments: **lumen**, where the four protein supercomplexes are embedded driving the electron transport in two modes, linear and cyclic; and the **stroma**, compartment of the C3 photosynthetic carbon fixation. The cytosol defines the system boundary. In colour (green and blue) we have highlighted the reactions linking the two submodels: the production and consumption of ATP and NADPH.

We included two compartments in our model, the thylakoid lumen and the chloroplast stroma. **Lumen.** The reaction kinetics for oxidised plastoquinone (PQ), oxidised plastocyanin (PC), oxidised ferrodoxin (Fd), lumenal protons (H) and non-phosphorylated antenna (light harvesting complexes) were taken from [14]. The four-state description of the quencher activity, based on the protonation of the PsbS protein and activity of the xanthophyll cycle, was taken from our mathematical model of non-photochemical quenching, initially developed to study short-term light memory in plants [17]. The previous description of ATP and NADPH consuming reactions is supplemented by the detailed description of the CBB Cycle. **Stroma.** Processes of the CBB Cycle have been implemented as in the mathematical model of the Calvin photosynthesis by Poolman *et al.* [10], based on the original work of Pettersson and Ryde-Pettersson [9]. The original model reproduces different carbon fixation rates and reaction activities at different concentrations of external orthophosphate, and includes the conversion of fixed carbon into either triose phosphates or sugar and starch. This model has been parametrised for CO_2_ saturating conditions and we kept the same assumption for all our analyses. The previous description of ATP synthesis is supplemented in our model with the new rate *v*_ATPsynthase_, which depends on the proton motive force built up by the PETC activity. Moreover, the stromal concentration of NADPH is dynamic.

### 2.2 Model compartments and units

The merged models were developed for different organisms ([14] for *C. reinhardtii*, [17] for *A. thaliana* and [10, 9] based on data for isolated spinach chloroplasts) and express the concentrations and rates in different units. To keep the original structure of the models, but provide consistency, we have kept the original units for each of the compartments and used a conversion factor (*p*_convf_, see Supplement) to convert quantities where needed. Thus, the concentrations of proteins and pool sizes inside the lumen are expressed as in previous models of the electron transport [14, 17] in mmol(mol Chl)^−1^, and the first order rates in mmol(mol Chl)^−1^s^−1^. Concentrations of metabolites and pools inside the stroma are expressed in mM, as in [9, 10]. To convert the concentration of ATP produced through the electron transport chain activity, expressed in mmol(mol Chl)^−1^, to mM, used to express concentrations in the stroma, we made several assumptions, as in our previous models of photosynthesis [15, 14, 17], which were originally derived from Laisk *et al.* [26]: i) chlorophyll content is assumed to be fixed and equal to 350 · 10^−6^ mol per m^2^ thylakoid membrane; ii) the volume of thylakoid stroma and lumen are 0.0112 l m^−2^ and 0.0014 l m^−2^, respectively. Thus, 1 mmol(mol Chl)^−1^ corresponds to 2.5 · 10^−4^M in the lumen and 3.2 · 10^−5^M in the stroma.

### 2.3 Computational analysis

The model has been implemented using the modelbase software, a console-based application written in Python, developed by us earlier this year [25]. Stoichiometry and parameters are provided in the Supplement and as a text file, to be found on our GitHub repository (www.github.com/QTB-HHU/photosynthesismodel). Moreover, we provide a Jupyter Notebook, that allows the user to repeat all the simulations leading to the production of the figures presented in this manuscript.

### 2.4 Reliability of the model

We have assembled the model of photosynthesis adapting previously validated and published mathematical models of two interdependent processes. We have used the same parameters as reported in the previous work and did not perform any further parameter fits (the full list of parameters in Supplement Tables S1-S5). We have monitored the evolution of several critical parameters to evaluate physiological plausibility of our computational results, including lumenal pH (kept under moderate light around 6), RuBisCO rate (in the order of magnitude of measured values) and the redox state of the PQ pool, used as an estimate of the overall redox state. Moreover, systematic steady state analysis of the model under different light conditions lead to plausible concentrations of CBB cycle intermediates and fluxes, as reported in the literature [9].

## 3 RESULTS AND DISCUSSION

We employed our merged model of photosynthesis and carbon fixation to perform a systematic supply-demand analysis of the coupled system. First, we have integrated the system for various constant light intensities until it reached steady state. Examples are provided in the Supplement (Figure S1). We observed reasonable stationary values of intermediates and fluxes for most of the light intensities. However, under very low light intensities (below 5 µmol m^−2^s^−1^), the phosphorylated CBB cycle intermediates dropped to zero, and ATP assumed the maximal concentration equalling the total pool of adenosine phosphates. Depending on the initial conditions, either a non-functioning state, characterised by zero carbon fixation rate, or a functioning state, characterised by a positive stationary flux, was reached. This observation of bistability constituted the starting point of our analysis of the tight supply-demand relationship.

In order to analyse this behaviour in more detail, we performed time course simulations, in which the light was dynamically switched from constant sufficient light (between 20 and 300 µmol m^−2^s^−1^), to a “dark phase” of 200 s duration with a light intensity of 5 µmol m^−2^s^−1^, back to high light, and observed the dynamics of the model variables. In Figure 2 we display the dynamics of the internal orthophosphate concentration, the sum of all three triose phosphate transporter (TPT) export rates and the RuBisCO rate (from top to bottom respectively) during such light-dark-light simulation. In agreement with the steady-state simulations, higher light intensities result in a higher overall flux during the initial light phase. Higher carbon fixation and export fluxes are accompanied by lower orthophsophate concentrations, which reflect higher levels of CBB cycle intermediates. In the dark phase, the non-functional state with zero carbon flux is approached. While rates decrease, orthophosphate increases, reflecting a depletion of the CBB intermediate pools. In the second light phase, only the simulated transitions to light intensities of 150 and 200 µmol m^−2^s^−1^could recover a functional state under the chosen conditions. For lower light intensities, apparently the CBB intermediate pool was depleted to a level, at which re-illumination fails to recover the CBB cycle activity. Obviously, this behaviour disagrees with everyday observations in nature (plant leaves recover from dark periods also under low light intensities). Nevertheless, the model is useful to generate novel insight. First, it illustrates that there exists a critical threshold of intermediate concentrations. If levels drop below this threshold, the cycle cannot be re-activated. Second, it explains the mechanisms leading to intermediate depletion. Under low light conditions insufficient energy supply results in reduced activity of ATP and NADPH dependent reactions in the carbon fixation cycle, leading to a reduced regeneration rate of Ribulose 1,5-bisphosphate (RuBP) from Ribulose-5-Phosphate (Ru5P). Simultaneously, the reversible (ATP independent) reactions remain active. Since triose phosphates are products of reversible reactions, these continue to be exchanged via the TPT export reactions with free phosphate, which leads to a depletion of the CBB cycle intermediates and a concomitant increase of the orthophosphate pool.

**FIGURE 2.**
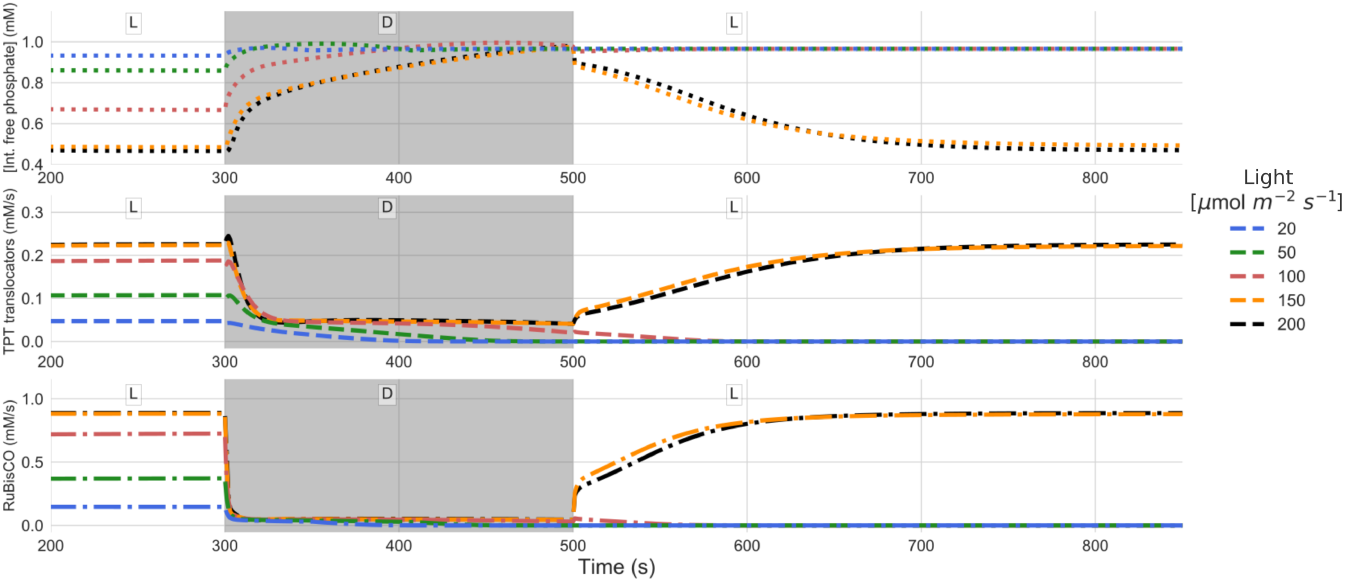
Simulations of light-dark-light transitions for different light intensities, ranging from 20-200 µmol m^−2^s^−1^. Shown are the dynamics of internal orthophosphate concentration, triose phosphate transporter (TPT) export and carbon fixation rates. The simulated time-courses are shown from 200s, when the system has reached a stationary state. From 300-500s (grey area), the external light has been set to 5 µmol m^−2^s^−1^. The figure illustrates that for low light intensities the CBB cycle fails to restart in the second light period.

Clearly, the model is missing important mechanisms that prevent such a functional failure. In particular, we are interested in how a **stand-by mode** can be realised, in which intermediate levels are maintained above the critical threshold, while at the same time the resources required to do so, are minimised. A possible strategy to prevent the collapse of the carbon fixation cycle is to resupply important intermediates. One biochemical process in plants that is known to produce Ru5P is the oxidative phase of the Pentose phosphate pathway (PPP), in which one Glucose-6-phosphate molecule is oxidised and decarboxylated to Ru5P, while producing NADPH and CO_2_ [27]. In order to estimate critical intermediate levels required to prevent the collapse of the carbon fixation cycle, we performed simulations under sufficient light (500 µmol m^−2^s^−1^), with different initial conditions. The initial concentrations of all carbon fixation intermediates are set to zero, except for Ru5P. The simulated Ru5P concentration, depicted in Figure 3, displays a characteristic dynamics. In the first seconds, the CBB cycle intermediates are equilibrated by the fast reversible reactions. If this concentration remains above the critical threshold of approximately 2.5 µM, the cycle reaches a functional state, if it falls below, it will collapse. Interestingly, the threshold concentration is rather independent of the light intensity (see Figure S2).

**FIGURE 3.**
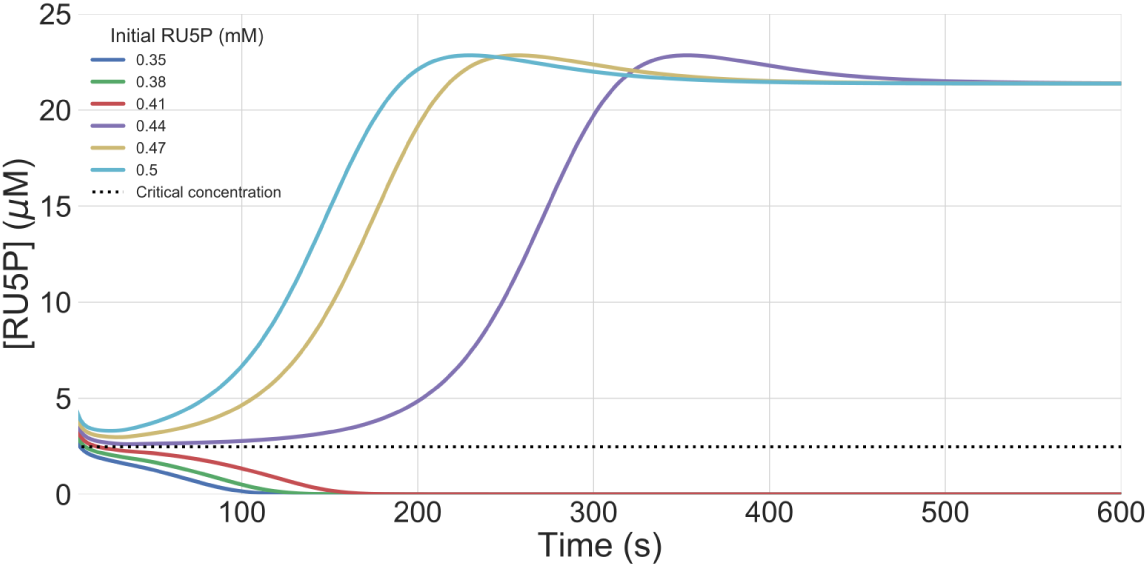
Simulations in light intensity of 500 µmol m^−2^s^−1^for different initial concentrations of RU5P, ranging from 0.35 to 0.5 mM. The RU5P abundance is shown after 10s, when the system is approximately equilibrated. The dashed line displays the critical concentration for sufficient cyclic activity after equilibrating. The figure displays that initial RU5P concentrations below 0.44 mM result in a loss of RU5P abundance.

To simulate a simple mechanism implementing a stand-by mode, which maintains sufficient CBB cycle intermediate levels, we introduced a trivial conceptual reaction, exchanging inorganic phosphate with Ru5P. Figure 4 displays simulated steady state values of the relative stromal ATP concentrations, Ru5P concentrations and lumenal pH in insufficient light conditions (5 µmol m^−2^s^−1^) as a function of the Ru5P influx. Again, a clear threshold behaviour can be observed. If the Ru5P influx exceeds approximately 4 µM/s, not only CBB intermediates assume non-zero concentrations, but also the lumenal pH reaches realistic and non-lethal levels. As expected, increased Ru5P influx results in its increased stationary concentrations, which is accompanied by an increased flux through RuBisCO and the TPT exporter (Figure S3), indicating a higher stand-by flux, and therefore, a higher requirement of resources to maintain this mode.

**FIGURE 4.**
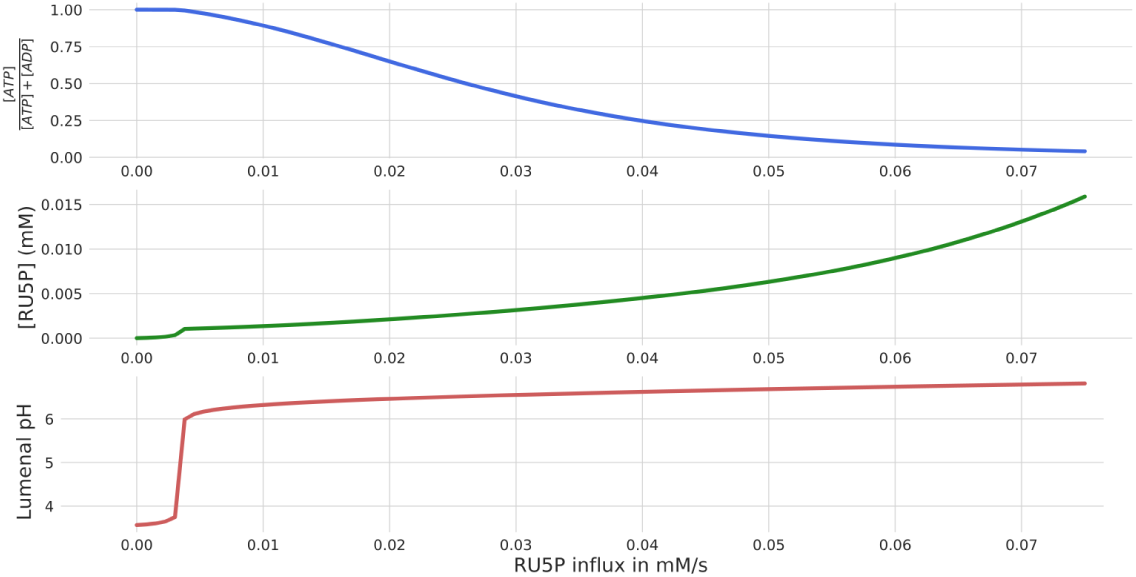
Steady state simulations in low light intensity of 5 µmol m^−2^s^−1^and systematically increasing influxes of Ru5P from 0. to 0.08 mM/s. The figure displays normalised ATP abundance, Ru5P concentration and lumenal pH.

These results suggest that a constant flux providing Ru5P in the dark with a rate just above the critical threshold of 4 µM/s should maintain intermediate CBB levels sufficiently high, while at the same time minimise the required investment. Indeed, with a constant supply of Ru5P with 5 µM/s, the system can be restarted and reaches a functional stationary state after a prolonged dark period (see Figure S4). Per carbon, this rate translates to 25-30 µM carbon/s, depending whether the pentoses are directly imported or derived from hexoses. Comparing this to stationary carbon fixation in the light of 0.1-1 mM/s (for light intensities between 20 and 200 µmol m^−2^s^−1^, see Figure 2 and Figure S1) shows that resupply under these conditions would consume a considerable fraction of the previously fixed carbon. This calculation demonstrates the importance of down-regulating the CBB cycle in dark conditions for a positive carbon fixation balance over a day/night cycle. Indeed, key enzymes in the carbon fixation cycle are known to be regulated by the pH and the redox state of the chloroplast stroma. For example, RuBisCO activity is controlled by proton levels and magnesium ions [28, 29]. Fructose-1,6-biphosphatase, Seduheptulose-1,7-biphosphatase and Phosphoribulokinase are all controlled by the redox state through the thioredoxin-ferredoxin system, and also by pH [30, 31, 32]. Furthermore, Hendriks *et al.* showed the light dependency of the ADP-glucose pyrophosphorylase [33], which is part of the lumped reaction *v*_Starch_ in our model. All these mechanisms will lead to a considerable reduction of the required stand-by flux of the CBB cycle, but are not yet included in our simple merged model.

In the original formulation of our model without constant Ru5P supply or light-dependent regulation of CBB enzymes, low light intensities lead to a rapid collapse of the cycle. In sufficient light, however, ATP levels are very high and carbon fixation rates are already saturated in moderate light conditions (Figure 2 and Figure S1). These findings indicate that the sets of parameters for the carbon fixation enzymes and the light reactions, derived from the respective original publications, might not be suitably adapted when employed in a merged, cooperating, system. This is not surprising considering that they originate from completely different systems and conditions.

In order to systematically investigate the supply-demand behaviour of the coupled system in different light conditions, we introduce a ‘regulation factor’ *f*_CBB_ of the CBB cycle, by which all *V*_max_-values of the light-regulated enzymes (see above) are multiplied. This allows for a systematic variation of the energy demand by simulating accelerated or decelerated carbon fixation activity. Performing this variation under different light conditions gives insight into the synchronisation of ATP and NADPH production and consumption rates, and thus enables a more profound analysis the supply-demand regulation of photosynthesis [34, 30, 31]. For the following steady-state analysis, the conceptual Ru5P influx reaction is not included. Figure 5 displays stationary values of key model variables for different light intensities and regulation factors. In agreement with the observations presented above that very low light intensities lead to a collapse of the cycle, ATP concentrations (Figure 5a) are maximal (zero ADP), triose phosphate export (Figure 5b) and starch production (Figure 5c) are zero, and the lumenal pH (Figure 5d) is very low (around 4). The latter is readily explained by the fact that the pH gradient built up by the low light cannot be reduced by the ATPase, which lacks the substrate ADP. Further, it becomes clear that the regulation factor of *f*_CBB_ = 1, corresponding to the original parameters, is far from optimal. The ATP:ADP ratio remains very high, and TPT export and starch production rates are well below their optimum, regardless of the light intensities. The stationary lumenal pH further illustrates that parameters are not ideally adjusted. Not only for very low light, but also for moderate to high light conditions (above 300 µmol m^−2^s^−1^) the lumen is dramatically acidic, indicating a mismatch in production and consumption processes. Increasing the regulation factor to values *f*_CBB ≈_ 4 leads to a dramatic improvement of the performance of the system. The ATP:ADP ratio assumes realistic and healthy values around one, triose phosphate export approximately doubles, and starch production increases by one order of magnitude compared to the original parameter values. Concomitantly, the lumenal pH remains moderate (>5.8, as suggested in [35]).

**FIGURE 5.**
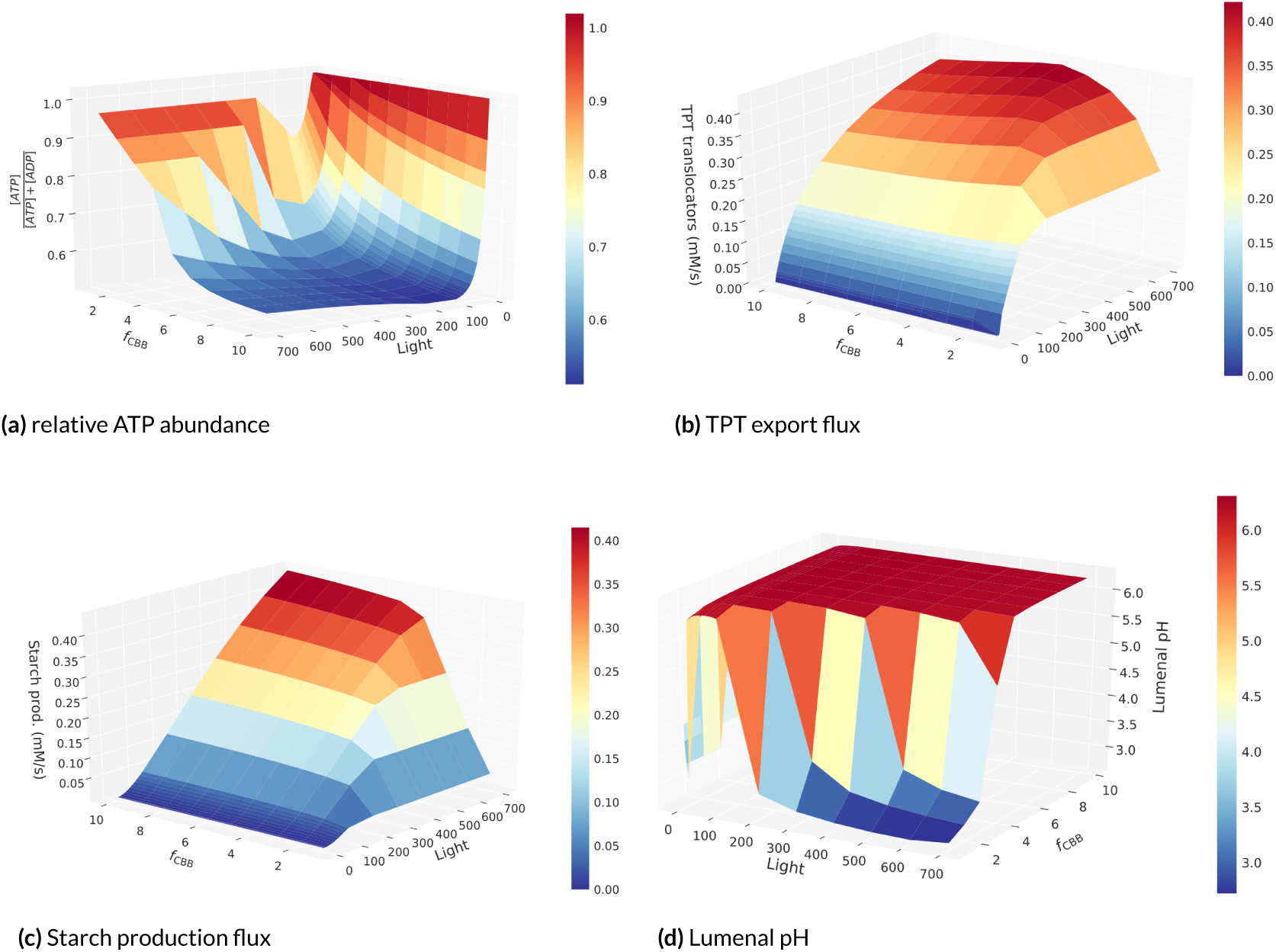
3D display of the steady state analysis of the system under varying light intensities (x-axis) and carbon fixation velocities (y-axis). On the z-axis: (a) the relative ATP abundance, (b) TPT export flux, (c) starch production rate, and (d) lumenal pH are displayed.

Quantitative analysis of the supply-demand behaviour of the system can be performed by calculating flux control coefficients [22, 23]. To investigate the relative overall flux control of supply and demand reactions, we first divide the set of all reactions in the model (R) into two non-overlapping sets S and D. S represents the supply set containing all PETC reactions and D represents the demand reaction set including all CBB cycle reactions. We define the the overall control of supply (*C*_Supply_) and demand (*C*_Demand_) reactions as the sum of the absolute values of all control coefficients of reactions from S and D, respectively, on the steady-state flux through the RuBisCO reaction,

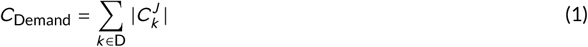

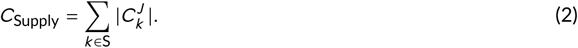

Figure 6 displays the normalized overall control of demand reactions *C*_Demand_/(*C*_Demand_ + *C*_Supply_), in dependence on different light intensities and carbon fixation regulation factors. Low light intensities and fast carbon fixation reactions shift the overall flux control to the supply reactions. This can readily be explained because under these conditions (low light and fast CBB enzymes) energy and redox provision by the light reactions are the limiting factor. Interestingly, PSII and PSI contribute strongest to the overall flux control on the supply side (Figure S5). Conversely, high light intensities and slow carbon fixation reactions shift the overall flux control to the demand side, because under these conditions, the system is energetically saturated, and the bottleneck is in the CBB cycle consuming the energy and redox equivalents. Noteworthy, it is the SBPase reaction that exhibits the highest overall flux control (Figure S6), while RuBisCO has only minor control.

**FIGURE 6.**
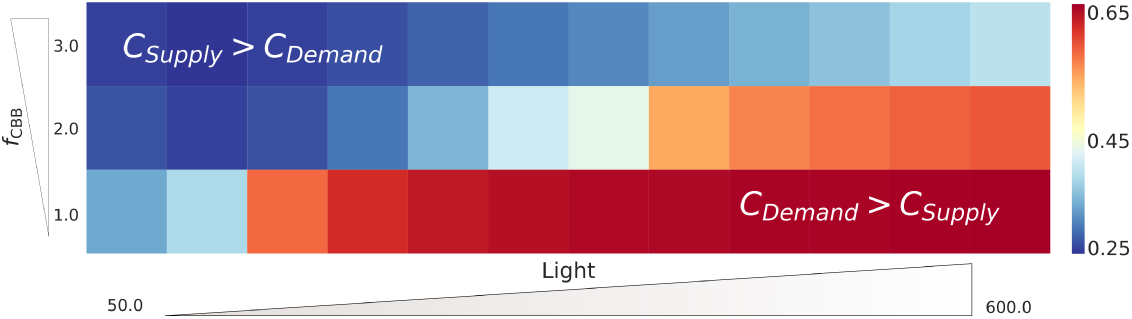
Normalized overall control of the demand reactions (*C*_Demand_) under different light intensities (x-axis) and CBB cycle activities (y-axis). The results show how the control shifts from the demand reactions under high light conditions, but low CBB activity, to the supply, under low light conditions but faster CBB cycle.

## 4 CONCLUSIONS

Merging mathematical models is a highly non-trivial task. Even if two individual models yield plausible results, there is no guarantee that this is also true after mathematically combining these models. Besides pure technicalities, such as converting concentrations to appropriate units, there are a number of issues that make merging models challenging. Commonly, individual models have been developed with quite different scientific questions in mind, and may therefore display drastically different degrees of details of the involved processes. Moreover, parametrisation is often performed for different organisms, tissues or conditions. Most importantly, increasing the system size by integrating two or more models may lead to novel emergent properties that were not observable in the individual models.

In this work, we have successfully merged a model of the PETC, supplying ATP and NADPH, to a model of the CBB cycle, consuming ATP and NADPH. The successful merge was largely facilitated by ensuring a comparable level of simplification of the two individual models (PETC described by 9 ODEs and CBB cycle by 15 ODEs). Our merged model represents a supply-demand system and as such exhibits systemic properties that did not exist in each of the individual models. Linking supply and demand processes into one functional model allowed us to employ metabolic control analysis for a systematic investigation of the regulatory dependence between the PETC and CBB cycle. By simulating the light-dark-light transitions we could rationalise the importance of the oxidative PPP in providing substrates as a mechanism to operate the CBB cycle in a stand-by mode. Simultaneously, we illustrate that regulating the activity of the CBB cycle in very low light is critical to avoid excessive investment into the stand-by mode. Moreover, the model demonstrates that regulation adapting to different light intensities is important to balance the supply by the PETC to the downstream demand. Using MCA, we quantified the control distribution of supply and demand in the system for different light conditions and for varying CBB cycle activities. By introducing a regulation factor, corresponding to the CBB cycle enzyme activities, we demonstrate that the system requires higher input of light to obtain saturation for faster carbon fixation. Our MCA analysis showed that supply reactions exhibit high overall flux control when the light is limited. Conversely, the demand reactions control the flux in light-saturating conditions. Among the supply reactions, the activity of PSII and PSI exhibit the highest overall flux control, while among the demand reactions, the SBPase maintains the highest overall flux control (see Figure S5 and Figure S6). Interestingly, the often considered bottleneck enzyme RuBisCO exhibits only little overall flux control. This observation can be explained by the fact that the model assumes saturated CO_2_ conditions.

Our model is freely available as open source software, and we ensure that the results presented here can easily be reproduced. Because of its balanced simplicity and clear modular structure, we envisage that it serves as a platform for future development. Only relatively minor modifications will be necessary to employ it for further analyses of the relationship between the PETC and other subprocesses. For instance, by describing starch as a dynamic variable and by providing a simplified representation of the oxidative PPP, one could improve our understanding of the light dependent turnover of starch [36] and rationalise the resupply of pentoses from hexoses in the chloroplast by the oxidative PPP [37, 38] and investigate the role of alternative shunts [39]. Moreover, by including a simplified representation of the photorespiratory pathway one could further investigate the energy balance [40] and the distribution of flux control between the PETC, the CBB cycle and the photorespiration reactions.

The process of integrating two models described here illustrates the strength of theoretical approaches. Linking two processes leads to novel properties (here supply-demand balancing), which can be investigated to provide new fundamental insight. The merged model can rationalise the importance of systemic properties, and thus explain *why* certain mechanisms exist. In particular, none of the individual models could have explained the relevance of the stand-by mode or the role of adaptive regulation in maximising efficiency, and thus explain the functional importance of the oxidative PPP or the redox and pH sensitivity of key CBB enzymes in a dynamic environment.

## A SUPPLEMENT

We are providing the mathematical description of the model of photosynthesis implemented in Python using the modelbase software developed in our Lab [25]. The code can be downloaded from the GitHub repository (https://github.com/QTB-HHU/photosynthesismodel) and is accompanied by a Jupyter Notebook that allows user to easily repeat the simulations included in the main text of the “Balancing energy supply during photosynthesis - a theoretical perspective” article.

### A.1 Mathematical Model

The mathematical model of photosynthesis is a result of merging previously developed mathematical models of the photosynthetic electron transport chain [14], complemented with the four-state description of the non-photochemical quenching [17], with the Poolman [10] implementation of the model of C3 photosynthesis carbon fixation [9].

#### A.1.1 Stoichiometry

The system of equations comprises a set of 24 coupled ordinary differential equations and 34 reaction rates. The first 9 differential equations describe dynamics of the PETC and follow the change of the oxidised fraction of the plastoquinone pool (Eq. S1), oxidised fraction of the plastocyanin pool (Eq. S2), oxidised fraction of the ferredoxin pool (Eq. S3), stromal concentration of ATP (Eq. S4), stromal concentration of NADPH (Eq. S5), proton concentration in the lumen (Eq. S6), phosphorylated fraction of light harvesting complexes (Eq. S7), the fraction of non-protonated PsbS protein (Eq. S8), and the fraction of violaxanthin in the total pool of xanthophylls (Eq. S9):

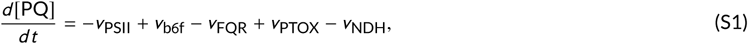

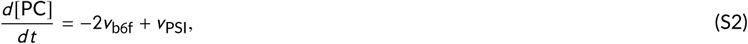

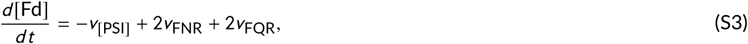

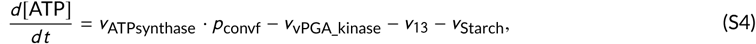

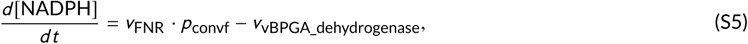

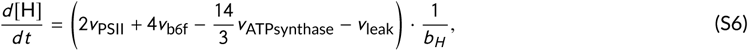

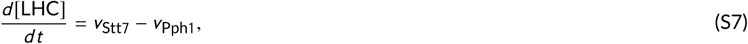

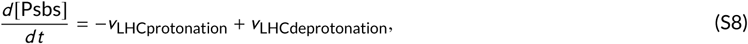

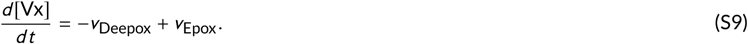

The next 15 differential equations govern the temporal evolution of the CBB cycle intermediates. The equations originate from the Pettersson and Ryde-Petterson model [9]:

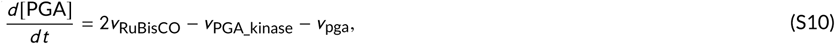

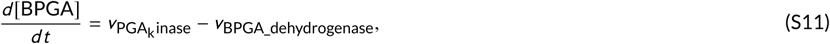

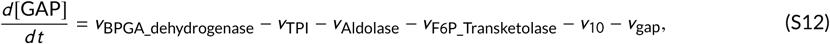

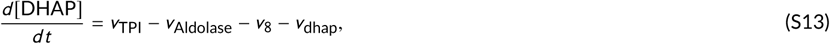

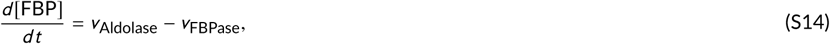

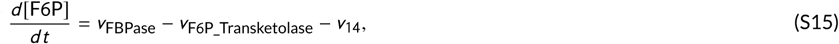

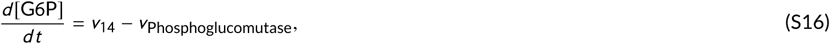

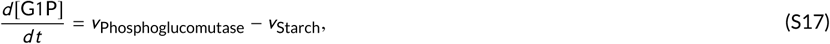

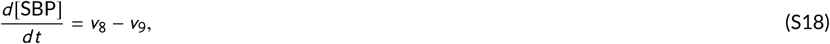

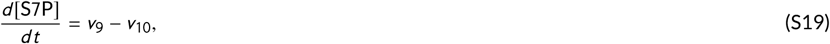

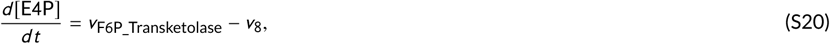

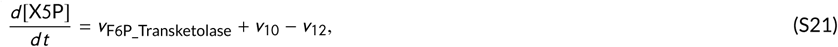

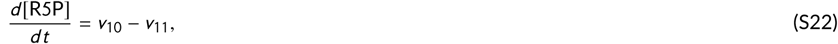

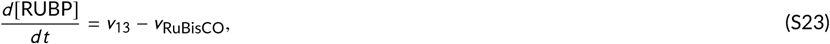

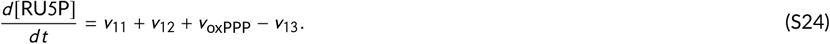

Moreover, using the functionality of modelbase [25], we have incorporated seven algebraic expressions from which we have derived seven dependent variables: the reduced fraction of PQ (Eq. S25), the reduced fraction of PC (Eq. S26), the reduced fraction of Fd (Eq. S27), the stromal concentration of ADP (Eq. S28), the stromal concentration of NADP (Eq. S29), the stromal concentration of orthophosphate (Eq. S30), and the inhibition factor of the triose phosphate translocators (Eq. S31):

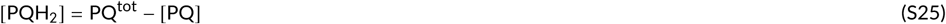

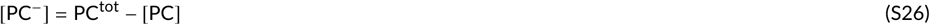

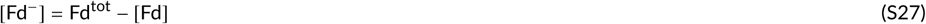

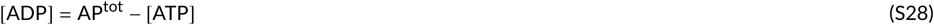

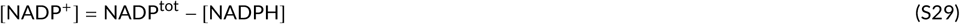

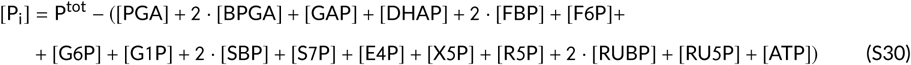

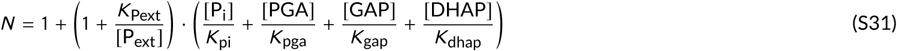

where P_ext_ denotes external orthophosphate and *K*_x_ represents the apparent dissociation constant for the formed complex between.

Additionally, the overall quencher activity can be derived using the following equation

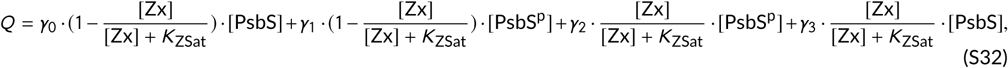

where [Zx] is the concentration of deepoxidised xantophylls ([X^tot^] = [Zx] + [Vx]), [PsbS^p^] is the concentration of protonated PsbS protein and the parameters are described in Table S4.

#### A.1.2 Reaction rates

The rate equations of all, except three reaction rates linking the submodels, have been kept as in the original works [9, 10, 14, 17]. We have substituted the two consuming reactions from [14] with the whole CBB cycle model and the one ATP synthase reaction of the [9] with the ATP synthase reaction from [14]. Nevertheless, to allow for an easy and self-sufficient read and to foster faster reproducibility of this work, we are providing below the full set of the reaction rates used for this model.

**The quasi-steady state approximation used to calculate the rate of photosystem II (PSII)**

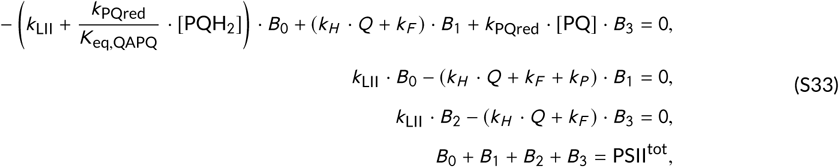

where *k*_LII_ is the light activation rate of PSII and *B_i_*, where *i ∈* (0, 1, 2, 3), is a temporary state of photosystem II, relating light harvesting capability with the occupation of reaction centres.

**The quasi-steady state approximation used to calculate the rate of photosystem I (PSI)**

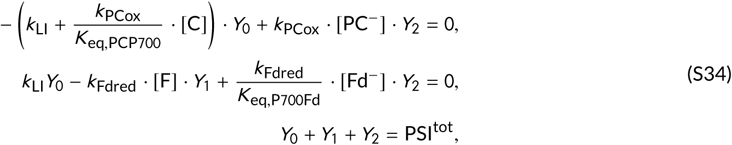

where *k*_LI_ is the light activation rate of PSI (determined by the total light intensity and the relative cross-section of PSI) and Y_*i*_, where *i ∈* (0, 1, 2) corresponds to one of the states of the PSI.

**The reactions of the PETC**

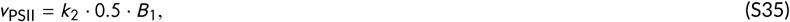

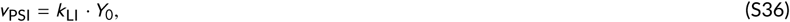

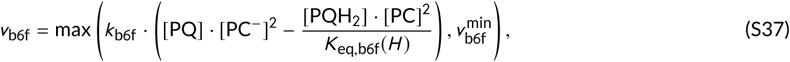

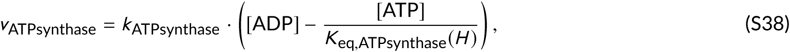

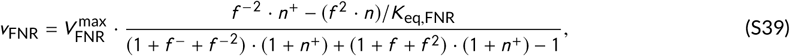

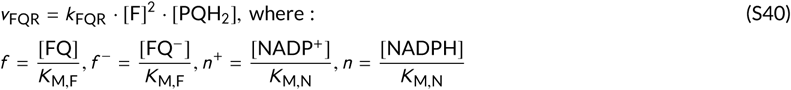

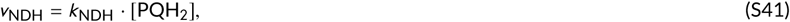

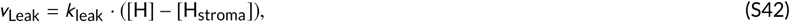

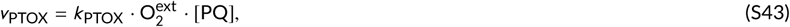

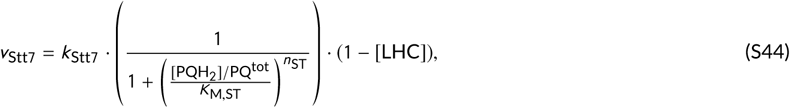

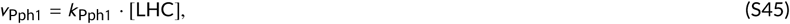

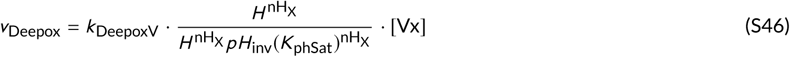

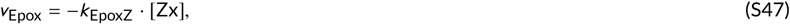

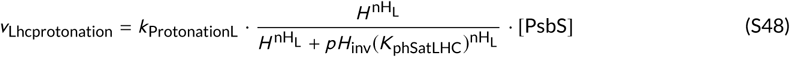

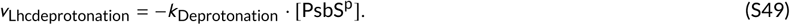

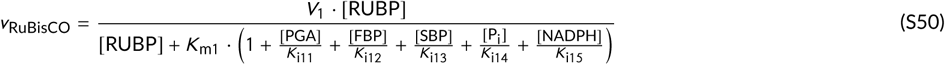

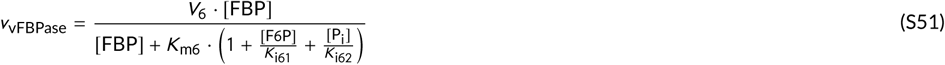

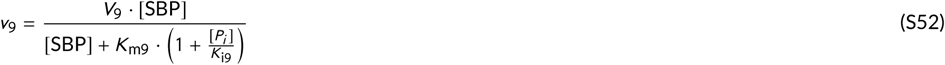

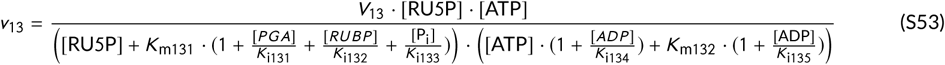

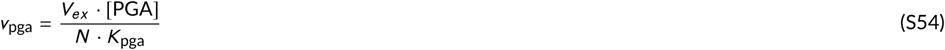

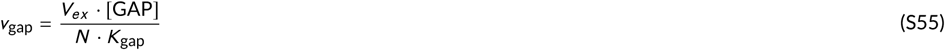

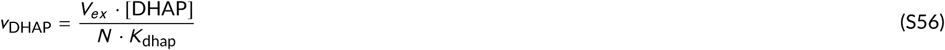

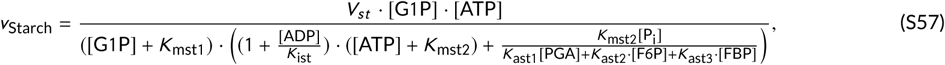

where the close-to-equilibrium approximations from the Pettersson and Ryde-Pettersson original model [9] has been dropped in favour of Poolman implementation [10], using fast rate constant (*k*):

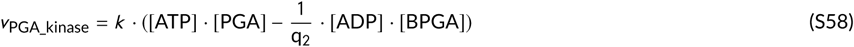

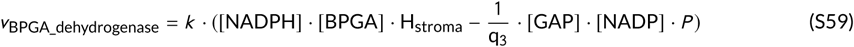

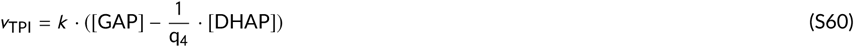

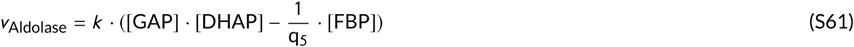

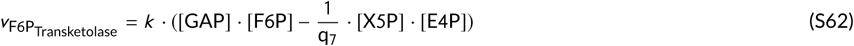

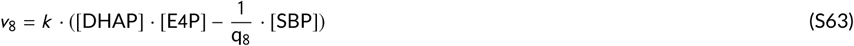

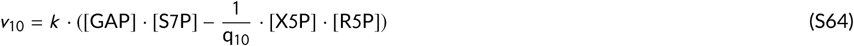

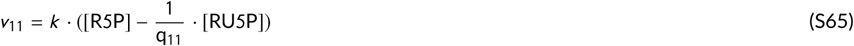

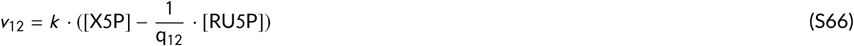

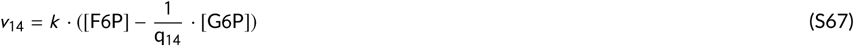

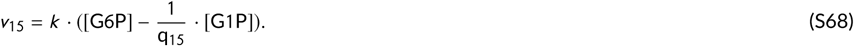

#### A.1.3 Parameters

The complete summary of parameters used in the model is included in Tables S1-S5.

**TABLE S1.** Pool sizes.

| Parameter | Value | Description and reference |
| --- | --- | --- |
| Luminal side [mmol(mol Chl)<sup>-1</sup>] |  |  |
| PSII<sup>tot</sup> | 2.5 | PSII reaction centres. Unchanged from [14] |
| PSI<sup>tot</sup> | 2.5 | PSI reaction centres. Unchanged from [14] |
| PQ<sup>tot</sup> | 17.5 | PQ + PQH<sub>2</sub>. Unchanged from [14] |
| PC<sup>tot</sup> | 4 | PC<sup>-</sup> + PC<sup>+</sup>. Unchanged from [14] |
| Fd<sup>tot</sup> | 5 | Fd<sup>-</sup> + Fd<sup>+</sup>. Unchanged from [14] |
| PsbS<sup>tot</sup> | 1 | relative pool of PsbS. Unchanged from [17] |
| X<sup>tot</sup> | 1 | relative pool of xanthophylls (Vx+Zx). Unchanged from [17] |
| O<sub>2</sub><sup>ex</sup> | 8 | external oxygen pool, corresponds to 250μM. Unchanged from [14] |
| Pi<sub>mol</sub> | 0.01 | internal pool of phosphates, required to calculate ATP equilibrium |
| Stromal side [mM] |  |  |
| AP<sup>tot</sup> | 2.55 | total adenosine phosphate pool (ATP + ADP). Increased, from Bionumbers |
| NADP<sup>tot</sup> | 0.8 | NADP + NADPH. Unchanged from [9] |
| P<sub>ext</sub> | 0.5 | external phosphate. Unchanged from [9] |
| CO<sub>2</sub> | 0.2 | Unchanged from [9] |

**TABLE S2.** Rate constants and key parameters of the PETC.

| Parameter | Value | Description and reference |
| --- | --- | --- |
| $k_{PQred}$ | $250 \text{ mmol}^{-1}(\text{mol Chl}) \text{ s}^{-1}$ | Unchanged from [14] |
| $k_{PCox}$ | $2500 \text{ mmol}^{-1}(\text{mol Chl}) \text{ s}^{-1}$ | Unchanged from [14] |
| $k_{Fdred}$ | $2.5 \cdot 10^5 \text{ mmol}^{-1}(\text{mol Chl}) \text{ s}^{-1}$ | Unchanged from [14] |
| $k_{Cytb6f}$ | $2.5 \text{ mmol}^{-2}(\text{mol Chl})^2 \text{ s}^{-1}$ | Unchanged from [14] |
| $k_{ATPsynthase}$ | $20 \text{ s}^{-1}$ | Unchanged from [14] |
| $k_H$ | $5 \cdot 10^9 \text{ s}^{-1}$ | rate of non-radiative decay. Unchanged from [14] |
| $k_F$ | $6.25 \cdot 10^8 \text{ s}^{-1}$ | rate of fluorescence. Unchanged from [14] |
| $k_P$ | $5 \cdot 10^9 \text{ s}^{-1}$ | rate of photochemistry. Unchanged from [14] |
| $k_{PTOX}$ | $0.01 \text{ mmol}^{-1}(\text{mol Chl}) \text{ s}^{-1}$ | Unchanged from [14] |
| $k_{NDH}$ | $0.004 \text{ s}^{-1}$ | Unchanged from [14] |
| $v_{b6f}^{min}$ | $-2.5 \text{ mmol}(\text{mol Chl})^{-1} \text{ s}^{-1}$ | Unchanged from [14] |
| $v_{FNR}^{max}$ | $1500 \text{ mmol}(\text{mol Chl})^{-1} \text{ s}^{-1}$ | Unchanged from [14] |
| $k_{FQR}$ | $1 \text{ mmol}^{-2}(\text{mol Chl})^2 \text{ s}^{-1}$ | Unchanged from [14] |
| $pH_{stroma}$ | 7.8 | stroma pH of a dark adapted state. Unchanged from [14] |
| $k_{leak}$ | $0.010 \text{ s}^{-1}$ | Unchanged from [14] |
| $b_H$ | 100 | Unchanged from [14] |
| HPR | $\frac{14}{3}$ | ratio of protons to ATP in ATP synthase. Unchanged from [14] |
| Michaelis constants |  |  |
| $K_{M,F}$ | $1.56 \text{ mmol}(\text{mol Chl})^{-1}$ | Unchanged from [14] |
| $K_{M,N}$ | $0.22 \text{ mmol}(\text{mol Chl})^{-1}$ | Unchanged from [14] |
| $K_{M,ST}$ | 0.2 | Unchanged from [14], to yield a PQ redox poise of $\approx 1:1$ |
| $K_{M,fdST}$ | 0.5 | Unchanged from [14] |

**TABLE S3.** Rate constants and key parameters of the Calvin Cycle.

| Parameter | Value | Description and reference |
| --- | --- | --- |
| **$V_{max}$ values of Calvin cycle enzymes** |  |  |
| $V_1$ | 2.72 mM s<sup>-1</sup> | |
| $V_6$ | 1.6 mM s<sup>-1</sup> | |
| $V_9$ | 0.32 mM s<sup>-1</sup> | |
| $V_{13}$ | 7.992 mM s<sup>-1</sup> | |
| $V_{st}$ | 0.32 mM s<sup>-1</sup> | |
| $V_x$ | 2 mM s<sup>-1</sup> | |
| **Equilibrium constants** |  |  |
| q2 | 3.1 * (10.0 ** (-4.0)) | Unchanged from [9] |
| q3 | 1.6 * (10.0 ** 7.0) | Unchanged from [9] |
| q4 | 22.0 | Unchanged from [9] |
| q5 | 7.1 mM<sup>-1</sup> | Unchanged from [9] |
| q7 | 0.084 | Unchanged from [9] |
| q8 | 13.0 mM<sup>-1</sup> | Unchanged from [9] |
| q10 | 0.85 | Unchanged from [9] |
| q11 | 0.4 | Unchanged from [9] |
| q12 | 0.67 | Unchanged from [9] |
| q14 | 2.3 | Unchanged from [9] |
| q15 | 0.058 | Unchanged from [9] |
| **Michaelis constants** |  |  |
| $K_{m1}$ | 0.02 mM | Unchanged from [9] |
| $K_{mCO2}$ | 0.0107 mM | Unchanged from [9] |
| $K_{m6}$ | 0.03 mM | Unchanged from [9] |
| $K_{m9}$ | 0.013 mM | Unchanged from [9] |
| $K_{m131}$ | 0.05 mM | Unchanged from [9] |
| $K_{m132}$ | 0.05 mM | Unchanged from [9] |
| $K_{m161}$ | 0.014 mM | Unchanged from [9] |
| $K_{m162}$ | 0.3 mM | Unchanged from [9] |
| $K_{mst1}$ | 0.08 mM | Unchanged from [9] |
| $K_{mst2}$ | 0.08 mM | Unchanged from [9] |
| $K_{pga}$ | 0.25 mM | Unchanged from [9] |
| $K_{gap}$ | 0.075 mM | Unchanged from [9] |
| $K_{dhap}$ | 0.077 mM | Unchanged from [9] |
| $K_{pi}$ | 0.63 mM | Unchanged from [9] |
| $K_{pxt}$ | 0.74 mM | Unchanged from [9] |
| $K_{i11}$ | 0.04 mM | Unchanged from [9] |
| $K_{i12}$ | 0.04 mM | Unchanged from [9] |
| $K_{i13}$ | 0.075 mM | Unchanged from [9] |
| $K_{i14}$ | 0.9 mM | Unchanged from [9] |
| $K_{i15}$ | 0.07 mM | Unchanged from [9] |
| $K_{i61}$ | 0.7 mM | Unchanged from [9] |
| $K_{i62}$ | 12.0 mM | Unchanged from [9] |
| $K_{i9}$ | 12.0 mM | Unchanged from [9] |
| $K_{i131}$ | 2.0 mM | Unchanged from [9] |
| $K_{i132}$ | 0.7 mM | Unchanged from [9] |
| $K_{i133}$ | 4.0 mM | Unchanged from [9] |
| $K_{i134}$ | 2.5 mM | Unchanged from [9] |
| $K_{i135}$ | 0.4 mM | Unchanged from [9] |
| $K_{ist}$ | 10.0 mM | Unchanged from [9] |
| $K_{ast1}$ | 0.1 | Unchanged from [9] |
| $K_{ast2}$ | 0.02 | Unchanged from [9] |
| $K_{ast3}$ | 0.02 | Unchanged from [9] |

**TABLE S4.**
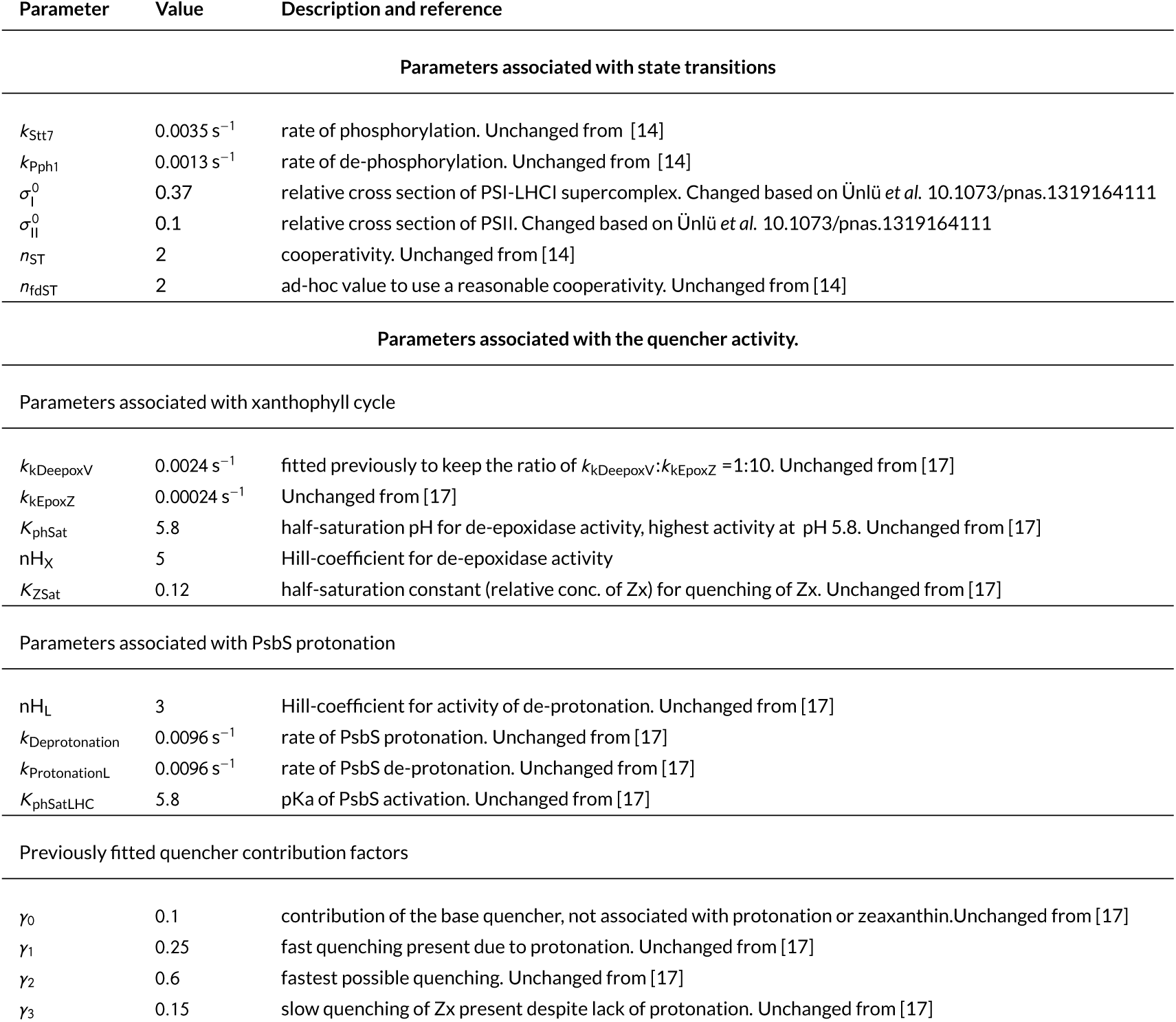
Parameters associated with photoprotective mechanisms.

**TABLE S5.**
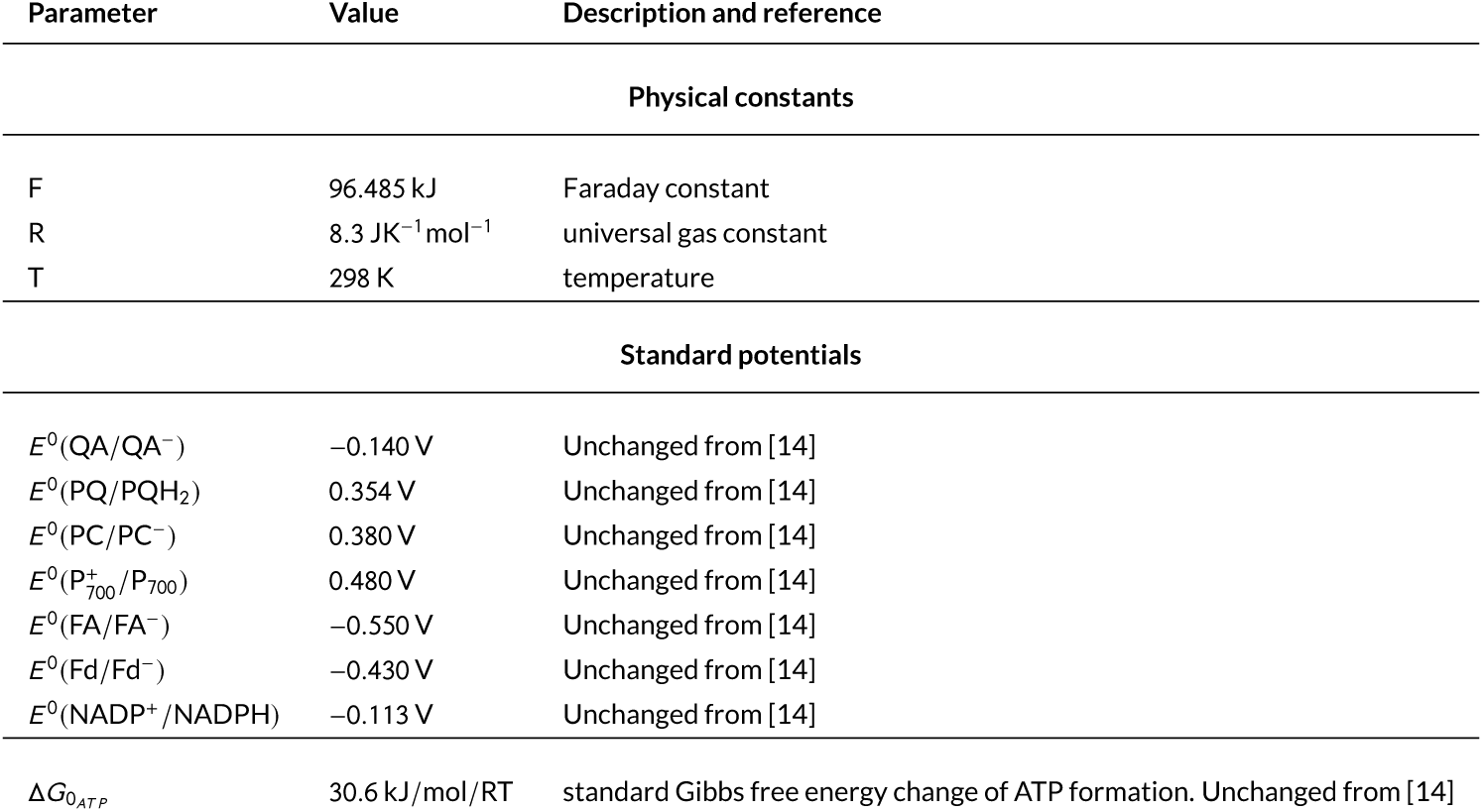
Physical constants and standard potentials

### A.2 Figures

Additional figures to support the Results section of the main text.

**FIGURE S1.**
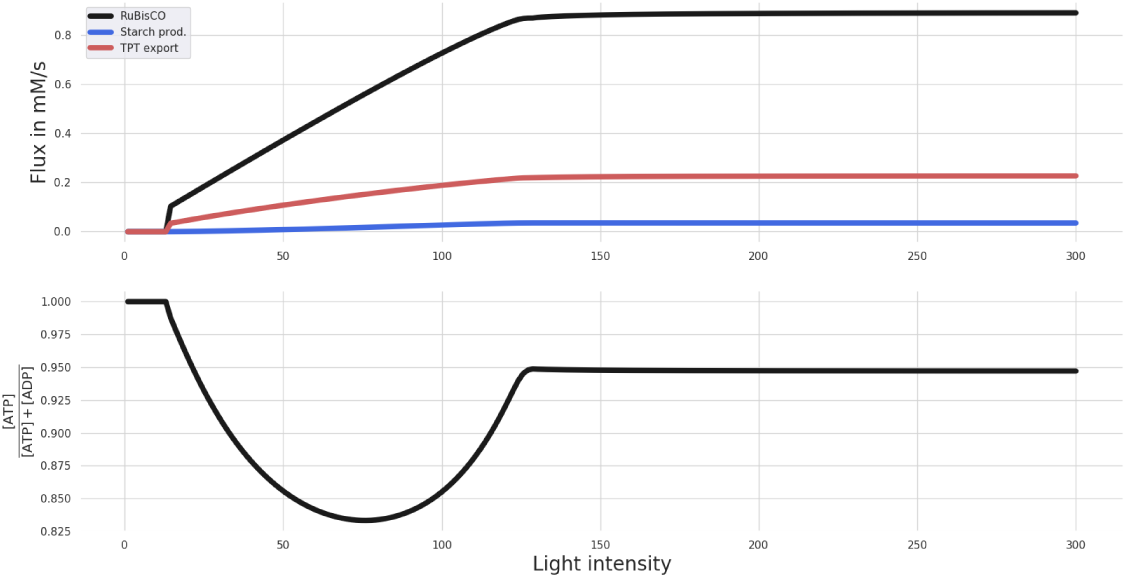
Result of the steady state simulation for different light intensities, varied between 0 and 300 µmol m^−2^s^−1^. Displayed are the rates of RuBisCO, starch production and TPT export (in mM/s).

**FIGURE S2.**
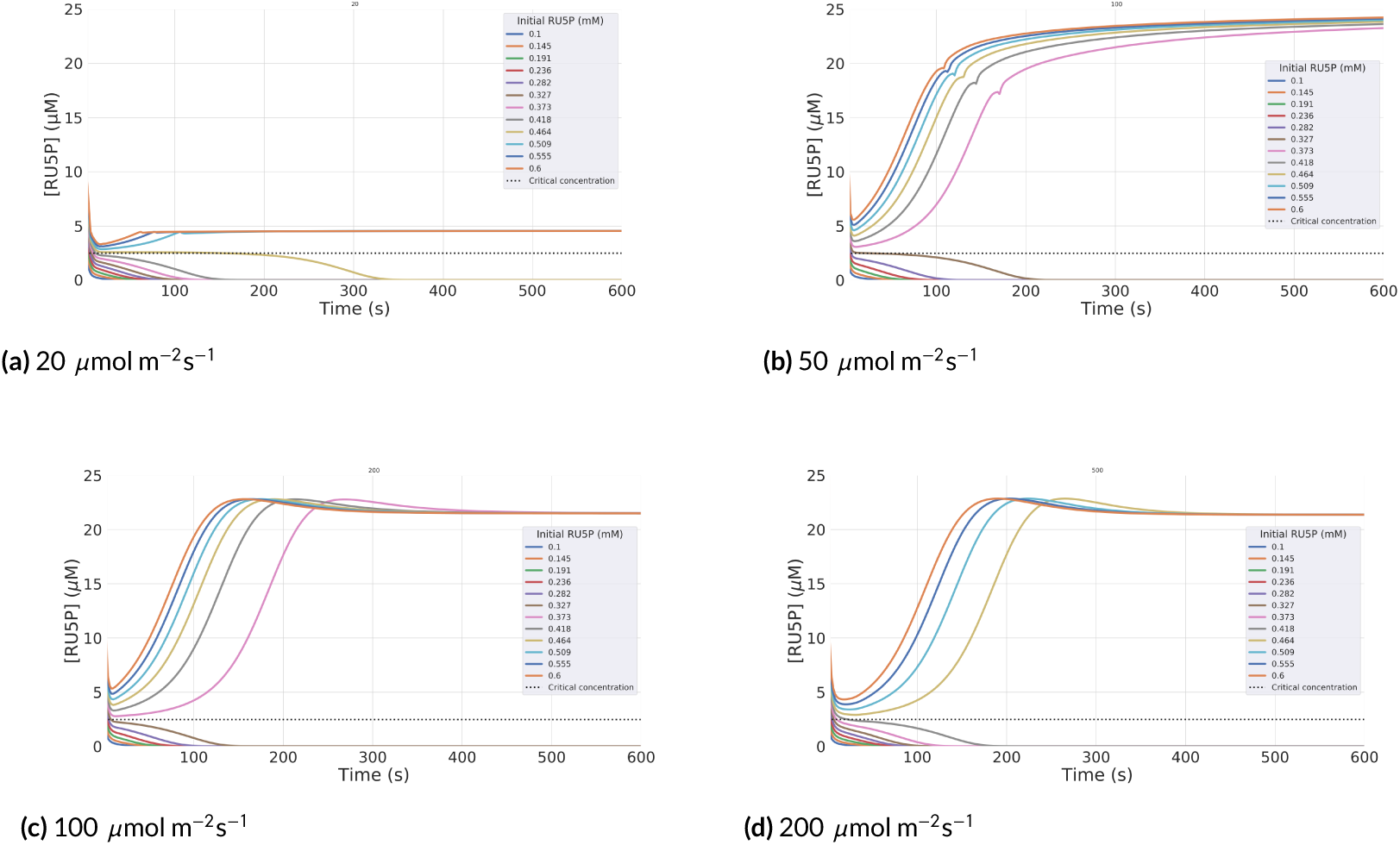
Simulations in different light intensities for different initial concentrations of RU5P, ranging from 0.1 to 0.6 mM. The RU5P abundance is displayed after 3 s, close to the beginning of equilibration. The dashed line displays the critical concentration of RU5P for sufficient cyclic activity after equilibrating. The critical concentration remains the same in different light intensities and energy abundance.

**FIGURE S3.**
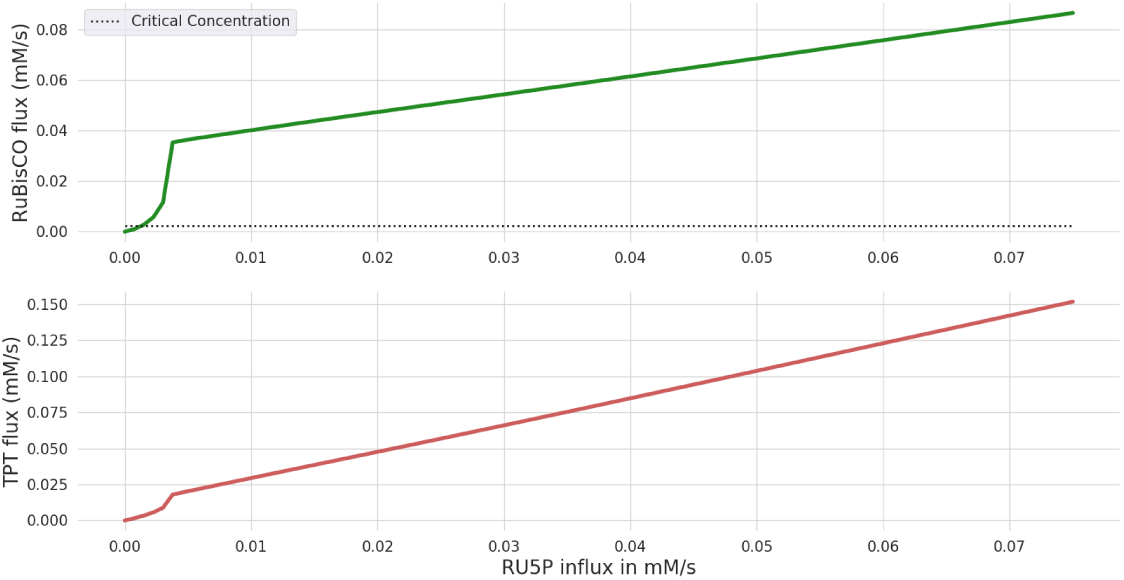
Steady state simulations in low light intensity of 5 µmol m^−2^s^−1^and systematically increasing influxes of RU5P from 0. to 0.08 mM/s. The figure displays rates of RuBisCO and triose phosphate export.

**FIGURE S4.**
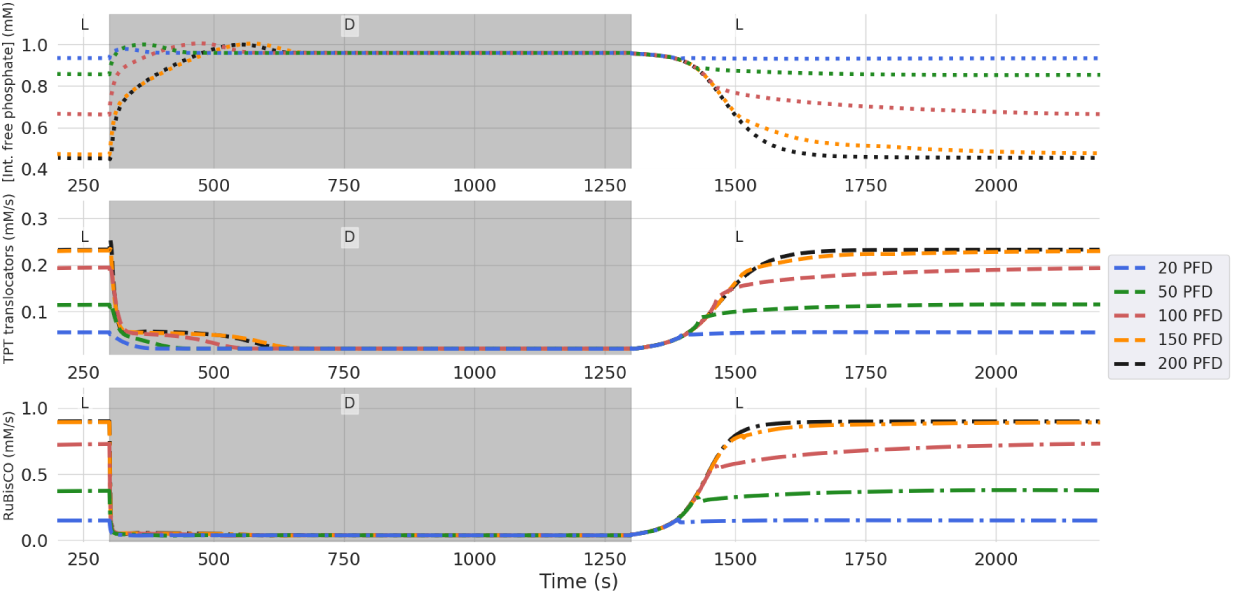
Simulations of light-dark-light transitions for different light intensities, ranging from 20-200 µmol m^−2^s^−1^with a constant RU5P influx of 5 µM/s. Shown are the dynamics of internal orthophosphate concentration, triose phosphate export and carbon fixation rates. The simulated time-courses are shown from 200s, when the system has reached a stationary state. From 300-1300s (grey area), the external light has been set to 5 µmol m^−2^s^−1^. The figure illustrates that even after long periods of low light intensities the CBB cycle is able to restart in the second light period if a small constant influx of RU5P is present.

**FIGURE S5.**
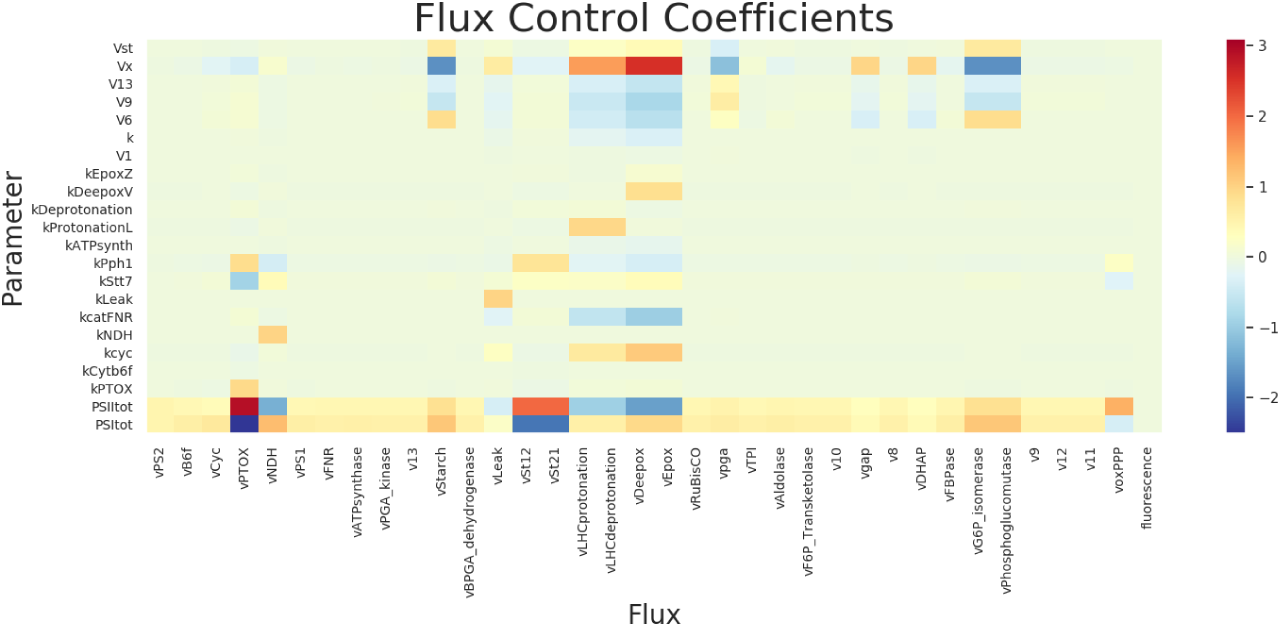
Flux control coefficients of the system in 50 µmol m^−2^s^−1^light and unchanged carbon fixation activity (*f*_CBB_ = 1). The parameters PSII^tot^ and PSI^tot^ denote the*V*_max_-values of Photosystem II and Photosystem I, respectively. They exhibit more flux control than other reactions.

**FIGURE S6.**
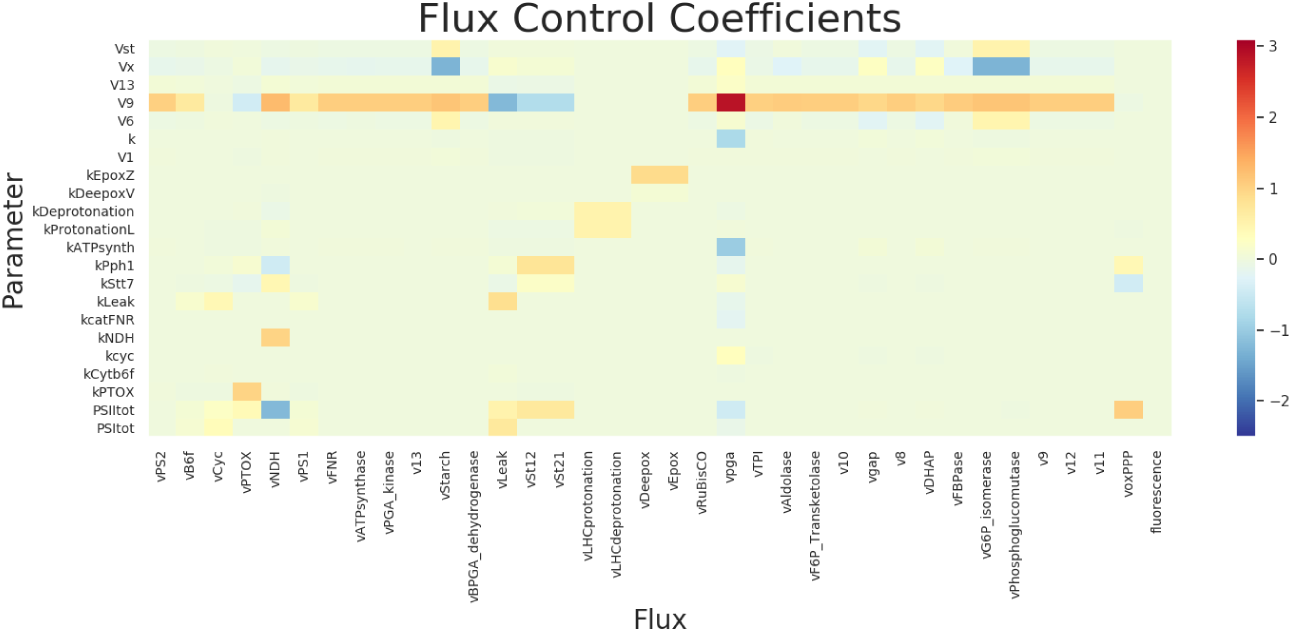
Flux control coefficients of the system in 300 µmol m^−2^s^−1^light and unchanged carbon fixation activity (*f*_CBB_ = 1). The parameter*V*_9_ denotes the*V*_max_-value of the SBPase and exhibits more flux control than other reactions.

## Abbreviations

CBB: Calvin-Benson-Bassham
Fd: ferrodoxin
MCA: Metabolic Control Analysis
NPQ: non-photochemical quenching
ODE: ordinary differential equations
PC: plastocyanin
PETC: photosynthetic electron transport chain
PFD: photon flux density
PPP: Pentose phosphate pathway
PQ: plastoquinone
PS: photosystem
RuBP: Ribulose 1,5-bisphosphate
Ru5P: Ribose-5-phosphate
TPT: triose phosphate transporters

